# Stathmin-2 Mediates Paracrine Hormone Regulation of Glucagon Through Lysosomal Trafficking in αTC1-6 cells

**DOI:** 10.64898/2026.04.02.715646

**Authors:** Nelson Chang, Samuel Ugulini, Savita Dhanvantari

**Author notes:** **Corresponding Author:** Savita Dhanvantari, PhD, Lawson Research Institute, 268 Grosvenor St, London ON N6A 4V2 CANADA. Equal first authors (Nelson Chang, Samuel Ugulini).

## Abstract

The secretion of glucagon from the pancreatic alpha (α) cell within the islets of Langerhans is physiologically regulated by nutrients (glucose, amino acids, fatty acids), neurotransmitters, and paracrine hormones. Insulin and somatostatin form an intra-islet paracrine network to control glucagon secretion through direct inhibitory effects on α cell secretory granule exocytosis. In a potential new cellular pathway for the regulation of glucagon secretion, we have previously identified the neuronal trafficking protein Stathmin-2 (Stmn2) as a negative regulator of glucagon trafficking and secretion by directing glucagon to degradative lysosomes. In this study, we examined if insulin and somatostatin direct glucagon to lysosomes in a Stmn2-dependent manner as part of their paracrine mechanisms. Using the αTC1-6 glucagon-secreting cell line and confocal microscopy of both fixed and live cells, we show that insulin and somatostatin direct glucagon, glucagon+LAMP1+ vesicles, and LAMP1-RFP to the intracellular region, away from sites of exocytosis. As visualized in live cells, insulin treatment resulted in the rapid retrograde transport of lysosomes from the cell periphery, and this effect was lost under siRNA-mediated silencing of Stmn2. Somatostatin appeared to enhance the intracellular retention of lysosomes, also in a Stmn2-dependent manner. We determined a possible mechanism for Stmn2 in the regulation of lysosome transport in αTC1-6 cells through the Arf-like small GTPase Arl8, indicating that Stmn2 may function in lysosomal positioning along microtubules. We propose that Stmn2-mediated lysosomal transport may be a potential new pathway, in addition to inhibition of secretory granule exocytosis, through which insulin and somatostatin regulate glucagon secretion.

## Introduction

The pancreatic alpha cell synthesizes and secretes glucagon, the major glucose counter-regulatory hormone. Under conditions of fasting, glucagon stimulates hepatic glycogenolysis and gluconeogenesis to maintain euglycemia. In addition to nutritional inputs from glucose, fatty acids and amino acids, and inputs from the central nervous system through neurotransmitters such as acetylcholine and norepineprhine, glucagon secretion is also regulated in a paracrine manner by intra-islet secretion of insulin and somatostatin and in an autocrine manner by glucagon ^1–3^.

Like all endocrine cells, the alpha cell stores its bioactive hormone glucagon in secretory granules until a stimulus triggers its release. This mechanism renders the alpha cell exquisitely sensitive to its microenvironment, and many studies have revealed how secretory granule exocytosis is regulated by intra-islet communication. In response to low glucose conditions (1-3 mM), glucagon acts in an autocrine feed-forward manner through glucagon receptor cAMP signalling in the alpha cell to maintain glucagon release during fasting ^4,5^. Insulin is secreted from the beta cell under high glucose conditions (> 7 mM), and is a very well-characterized and potent inhibitor of glucagon secretion. Through insulin receptor PI3 kinase signalling, both alpha cell membrane excitability and Ca^2+^ channel activity are reduced to suppress alpha cell exocytosis in the fed state ^6^. Additionally, the beta cell co-releases γ-amino butyric acid (GABA), which binds to GABAA receptors on the alpha cell and inhibits glucagon secretion through Cl^-^channel-induced membrane hyperpolarization ^7–9^. Somatostatin may control glucagon secretion in two ways: by fine-tuning the alpha cell response to low glucose ^10,11^, and by enhancing insulin-mediated inhibition at high glucose ^12^. At 3 mM glucose, delta cells release somatostatin to control glucagon secretion via SSTR2-cAMP-G_αi_ signalling ^13,14^ that may also reduce glucagon secretory granule docking to the plasma membrane for exocytosis. In the fed state, insulin induces somatostatin release from δ cells through gap junctions, further inhibiting glucagon secretion through alpha cell SSTR2 ^12,15^. Therefore, glucagon secretion is tightly regulated in response to nutritional cues through the interplay of intra-islet hormones.

In addition to regulation of alpha cell secretory granule exocytosis, there is emerging evidence that glucagon secretion may be also regulated by lysosomal trafficking, as is insulin ^16,17^. Under conditions of starvation, glucagon secretion may be regulated by crinophagy ^18^. Our lab has discovered stathmin-2, a neuronal protein associated with microtubule destabilization ^19^, plays a role in the trafficking of glucagon to intracellular lysosomes, presumably for degradation.

Overexpression of Stmn2 inhibits both basal and K^+^-stimulated glucagon secretion by transporting glucagon to lysosomes ^20^, while Stmn2 deficiency is associated with increased basal glucagon secretion in cells and in a model of uncontrolled Type 1 diabetes (T1D) and hyperglucagonemia ^20,21^. We have proposed that the negative regulation of glucagon through lysosomal trafficking is part of normal alpha cell homeostasis ^21,22^, and therefore such a pathway may be a novel means by which insulin and somatostatin regulate glucagon secretion. Therefore, we hypothesize that Stmn2 may mediate the paracrine control of glucagon secretion through directing glucagon to intracellular lysosomes for degradation.

In this study, we used the αTC1-6 cell line and immunofluorescence confocal microscopy to assess the trafficking of glucagon to lysosomes located in the intracellular region for degradation, or proximal to the plasma membrane for secretion, after treatment with either insulin or somatostatin. We show that insulin and somatostatin direct glucagon to intracellular lysosomes, and that this trafficking is dependent on Stmn2.

## Methods

### Cell line

Alpha TC1 clone 6 (αTC1-6) cells (RRID:CVCL_B036; a kind gift from C. Bruce Verchere, University of British Columbia, Vancouver, BC, Canada) were maintained in 25 mM glucose Dulbecco’s Modified Eagle Medium (DMEM) (Gibco^TM^, cat#11995073) supplemented with 15% horse serum (HS), 2.5% fetal bovine serum (FBS), and 100 units/ mL penicillin-streptomycin (Life Technologies, cat#15070063) at 37°C and 5% CO_2_ and 100% humidity.

When αTC1-6 cells reached 80% confluency, cells were washed with phosphate-buffered saline (PBS), lifted using 0.25% trypsin/EDTA (Thermo Fisher Scientific, cat#25200072), and resuspended in DMEM. Cells were seeded on sterile coverslips (#1.5, 0.17 mm thickness, 18 mm X 18 mm) in 6-well plates for all experiments, and cells were used until passage 12.

### Treatment with paracrine factors

αTC1-6 cells under passage six were grown to 60 to 70% confluency in 6-well plates and treated overnight with vehicle alone, 1 nM insulin plus 25 μM γ-amino butyric acid (GABA) or 1 nM insulin alone, or 400 nM somatostatin (Sigma-Aldrich, cat#S1763).

### Transfection

LAMP1-RFP was a gift from Walther Mothes (Addgene plasmid#1817; http://n2t.net/addgene:1817;RRID:Addgene_1817). Stmn2-GFP was obtained as the Stmn2 mouse tagged ORF clone (OriGene, cat#NM_025285) and pcDNA3-eGFP was used as the vector control. Cells were transiently transfected at 60-70% confluency in 6-well plates. For overexpression experiments, expression plasmids (Stmn2-GFP, LAMP1-RFP, or GFP-alone) were diluted in Opti-MEM reduced serum medium (Thermo Fisher Scientific, cat#31985062) and combined with Lipofectamine 2000 (Invitrogen, cat#1168019) in Opti-MEM medium for 5 minutes at room temperature before addition to serum-free media (Gibco^TM^. cat#11960044) supplemented with sodium pyruvate to induce uptake of plasmids. For gene silencing experiments, cells were transfected with either 40 nM pooled siRNAs targeting Stmn2 (Thermo Fisher Scientific, cat#4390771; s73354, s73355, s73356) or the siRNA negative control (Thermo Fisher Scientific, cat#4390846) with scrambled sequences that do not target any gene product, as done previously ^20,23^. Cells were transfected for 6 h, then incubated in regular culture medium overnight for cell recovery.

### Immunofluorescence staining

Cells were seeded on collagen-coated coverslips (100,000 cells per coverslip) and allowed to adhere overnight. After transfections and treatments, cells were washed in phosphate-buffered saline (PBS, pH 7.4), fixed with 2% paraformaldehyde (in PBS) for 30 min and permeabilized with 0.1% Triton X-100 (in PBS) for 5 min. After 2h incubation in blocking buffer (2% BSA + 0.05% Tween 20 in PBS) at room temperature (RT), cells were incubated overnight at 4°C with primary antibodies (diluted in blocking buffer) against the following proteins: **glucagon** (mouse anti-glucagon antibody, cat#10988, Abcam; 1:250); **Stmn2** (goat anti-SCG10 antibody, cat#115513, Abcam; 1:250); the lysosomal marker **LAMP1** (rabbit anti-LAMP1 antibody, cat#208943, Abcam; 1:100); the secretory granule exocytosis marker **Syntaxin1A** (rabbit anti-Syntaxin1A antibody, cat#272736, Abcam; 1:50); Transcription Factor EB (**TFEB)** (rabbit anti-TFEB antibody, cat#264421, Abcam; 1:50) or the lysosomal transport-associated small GTPase **Arl8A** (rabbit anti-Arl8A antibody, cat#17060-1-AP, Proteintech; 1:250). After washing in PBS, coverslips were incubated for 90 min at room temperature in the dark with the following fluorophore-conjugated secondary antibodies, as appropriate: Alexa Fluor^TM^ 594 donkey anti-mouse IgG antibody, cat#A21203, Thermo Fisher Scientific; 1:500); Alexa Fluor^TM^ 488 donkey anti-goat IgG antibody, cat#A11055, Thermo Fisher Scientific; 1:500); or Alexa Fluor^TM^ 488 donkey anti-rabbit IgG antibody, cat#A21206, Thermo Fisher Scientific; 1:500). Coverslips were then washed with PBS, DAPI (1:1000 in PBS) was added, and coverslips were mounted on microscope slides with anti-fade medium (Invitrogen, cat#2527959), sealed with nail polish, and stored at 4°C until analysis.

### Image acquisition

For fixed cells, imaging was done on a Nikon A1R confocal microscope equipped with NIS-Elements AR 5.41.01 software, using 60X oil-immersion objective with Galvano laser, at 1024 x 1024, pinhole size (1), average (16), and dwell time (2.4). The microscope was equipped with four lasers, including DAPI (405 nm), FITC (490 nm-519 nm), TRITC (561-594 nm), and Cy5 (633-647 nm). Images were processed using NIS Elements AI algorithm (restore.ai) to restore lost signals and then 2D deconvolution for maximum contrast and brightness. For live cell imaging, images were captured before, during and five minutes after 1 nM insulin or 400 nM somatostatin treatment. Imaging parameters were the same as for fixed cells except for average (1) and dwell time (1.1), sacrificing resolution in exchange for minimal rendering delays and real-time capture of signals over time.

### Image analysis

#### Cellular region definition

Adapted from a previously described method used for analysis of endosome and lysosome distribution ^24–26^, cellular regions of αTC1-6 cells and islets were defined using the fluorescence intensity of different known cellular markers along a distance from the center of the nucleus to the border of the cell periphery. Specifically, DAPI, LAMP1 and Syntaxin1A fluorescence were used to define the nuclear region, the intracellular region containing lysosomes and the cell periphery where exocytosis occurs, respectively. Lines were manually drawn from the center of the nucleus to the cell membrane in a manner that captured signals from all regions. The intensity of signals was quantified along the distance of the line on ImageJ Fiji. The intensity across the distance was then normalized to each individual cell’s nuclear radius to account for differences in cell shape and size. In this manner, the plot profile for LAMP1 was defined as the “intracellular region”, and that for Syntaxin1A, an exocytosis marker, was defined as the “cell periphery” region.

#### Co-localization of proteins and re-distribution analysis

Images from individual channels were processed using the “Image Calculator” function on ImageJ Fiji under the “AND” operation so that only the pixels containing two wavelengths would be captured with adjusted intensity. Plot profiles of the co-localized pixels were generated, and the re-distribution of the co-localized signals was quantified and categorized to the “intracellular region” or the “cell periphery” region, as described above.

### Statistical analysis

Individual coverslips were considered biological replicates for all experiments. Technical replicates for microscopy included multiple fields of view per coverslip (2-4). Plot profiles were used to quantify the fluorescence intensity of different proteins along a distance within an individual cell, then a two-way ANOVA analysis was conducted with variables including paracrine treatment and protein localization. The nuclear translocation of TFEB was calculated as the fluorescence intensity of TFEB within the nucleus stained by DAPI, normalized to the cytoplasmic fluorescence intensity of TFEB. A direct comparison of transcription factor EB fluorescence intensity was conducted using a two-sample t-test. Significance for all statistical analyses was set at α ≤ 0.05.

## Results

### Paracrine factors redistribute glucagon-positive vesicles from the cell periphery to the intracellular region

After treatment of αTC1-6 cells with 1 nM insulin/25 μM GABA or 400 nM somatostatin, the spatial localization of fluorescence intensities of the colocalized glucagon/Stmn2 (glucagon in secretory granules and lysosomes), glucagon/LAMP1 (glucagon in lysosomes) or glucagon/Syntaxin1A (glucagon at sites of exocytosis) were analyzed by plot profiles.

Glucagon/Stmn2 colocalized signals showed the strongest signal intensity near the plasma membrane (**Figure 1a, top row**). Upon treatment with 1 nM insulin/25 μM GABA or 400 nM somatostatin, the co-localized signals redistributed intracellularly, and the signals in the periphery were no longer as prominent (yellow arrowheads). Plot profile analysis showed a significant reduction in colocalized fluorescence intensity in the cell periphery upon treatment with either insulin/GABA (p<0.0001) or somatostatin (p<0.0001) **(Figure 1b, left panel)**. In addition, there was significantly more signal in the intracellular region compared to the periphery within the insulin/GABA-treated (p<0.001) and somatostatin-treated (p<0.001) groups (**Figure 1b**, left panel).

**Figure 1.**
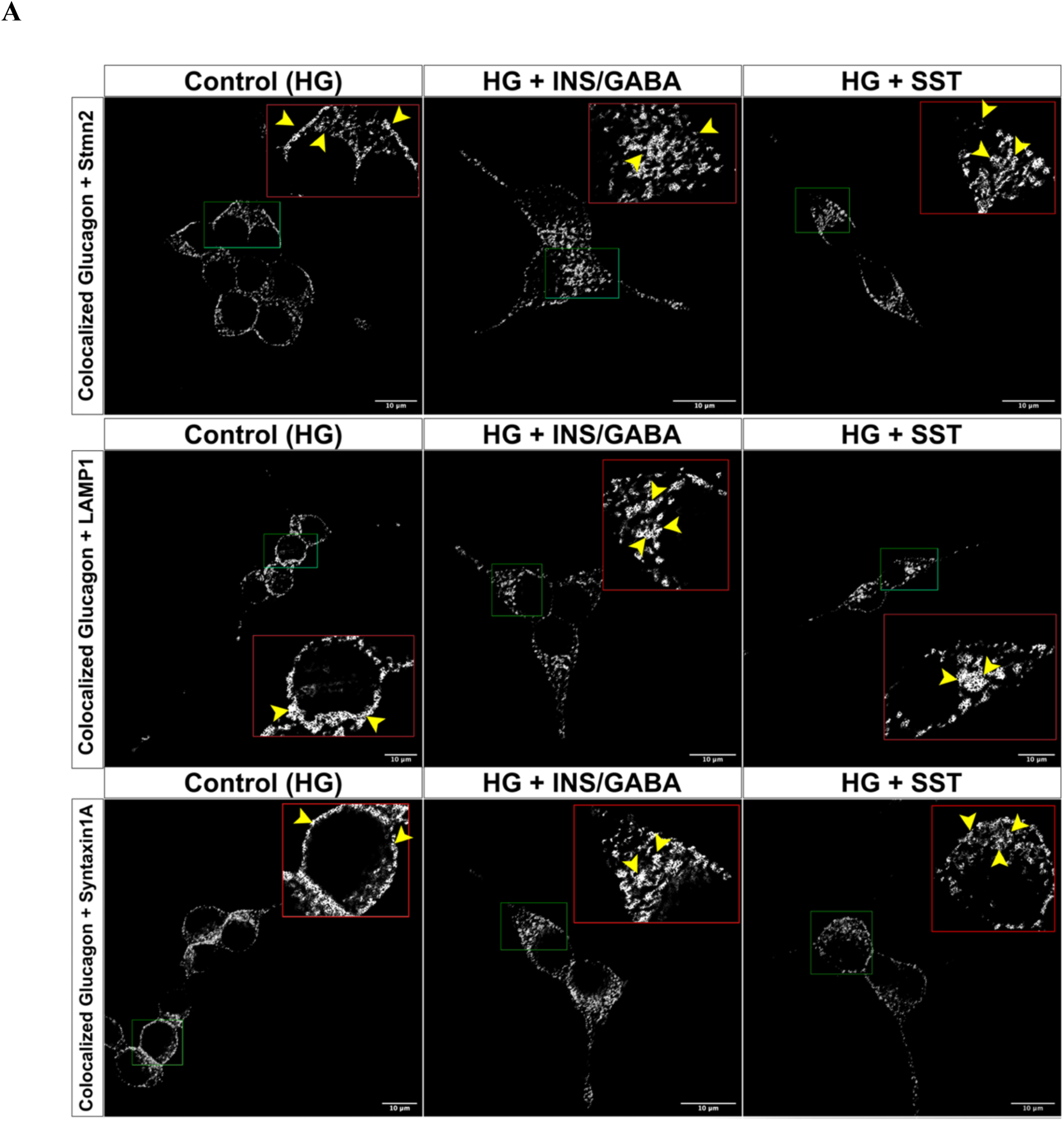
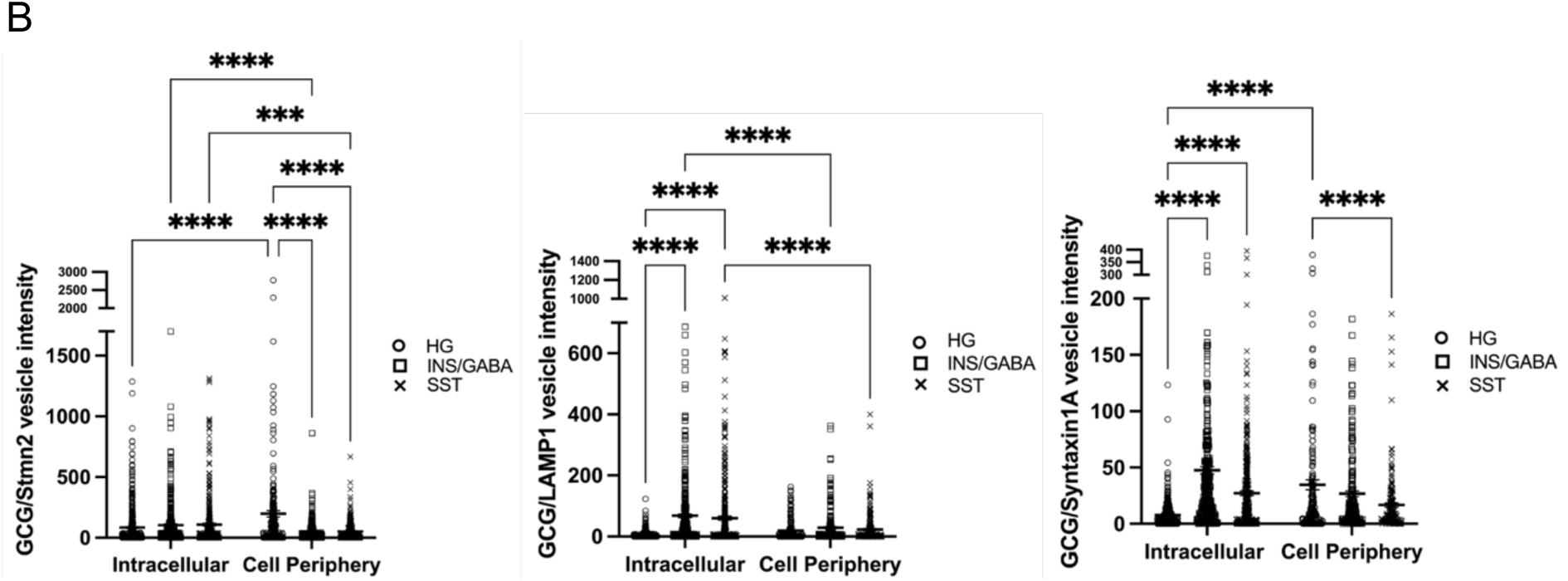
Redistribution of glucagon-containing vesicles to the intracellular region in αTC1-6 cells in response to the paracrine factors insulin/GABA or somatostatin (SST). A) Images show co-localized immunofluorescence signals for glucagon and Stmn2 (top row), glucagon and the lysosomal marker LAMP1 (middle row) and glucagon and the secretory granule membrane docking protein syntaxin-1A (bottom row). Insets show magnified areas outlined in green. Yellow arrowheads indicate the spatial localization of immunofluorescence signals in peripheral and intracellular regions. B) Plot profile analyses of colocalized immunofluorescence signals in the intracellular or peripheral regions in response to vehicle (open circle), 1 nM insulin+25 μM GABA (open squares), or 400 nM SST (x). Bars represent the averaged colocalized fluorescence intensities ± SEM (n=3) and symbols represent measurements from individual cells. Comparisons were made between and within cellular regions using a two-way ANOVA, followed by a post-hoc test. ***p<0.001, ****p<0.0001.

The glucagon/LAMP1 colocalized signal after treatment with 1 nM insulin/25 μM GABA or 400 nM SST revealed the paracrine-induced distribution of glucagon contained in lysosomes (**Figure 1a, middle row)**. The glucagon/LAMP1 colocalized signal was the most prominent near the plasma membrane and redistributed to the intracellular region upon treatment with insulin/GABA or somatostatin (yellow arrowheads). Plot profile analysis showed a significant increase (p<0.0001) in the colocalized signal in the intracellular region in response to 1 nM insulin/25 μM GABA or 400 nM somatostatin **(Figure 1b, middle panel)**.

The glucagon/Syntaxin1A colocalized signal in the presence of 1 nM insulin/25 μM GABA or 400 nM SST reveals the paracrine-induced distribution of glucagon inside secretory granules **Figure 1a, bottom row)**. The colocalized glucagon/Syntaxin 1A signal was the most prominent near the plasma membrane and redistributed to the intracellular region upon treatment of insulin/GABA or somatostatin (yellow arrowheads). Plot profile analysis showed the glucagon/Syntaxin1A colocalized signal in the intracellular region increased significantly (p<0.0001) in response to either 1 nM insulin/25 μM GABA or 400 nM somatostatin (**Figure 1b, right panel)**. In addition, peripheral colocalized signal was significantly reduced (p<0.001) by 400 nM somatostatin.

### Insulin and somatostatin mediate the intracellular trafficking of glucagon in a Stmn2-dependent manner

We then examined how manipulation of Stmn2 expression affected the subcellular distribution of glucagon in αTC1-6 cells, and if the effects of insulin and somatostatin may be mediated by Stmn2. Transfection of GFP alone was the negative control for transfection. In **Figure 2a**, **b** and **c,** the channels showed the relative fluorescence intensity of Stmn2 across treatments, with Stmn2-KD showing the least amount of fluorescence signal, and Stmn2-OE showed the strongest signal while distributing mainly in the intracellular region. Quantitatively, the integrated fluorescence intensity of Stmn2 after knockdown was significantly lower than the control (p<0.0001). In contrast, Stmn2-GFP fluorescence in the overexpression model was significantly higher than the control (p<0.01) **(Figure 2d)**. Under control conditions, glucagon immunofluorescence was largely distributed to the cell periphery (**Figure 2b**, yellow arrowheads) with some immunofluorescence in the intracellular region (quantified in **Figure 2e**). After Stmn2 knockdown **(Figure 2b)**, glucagon immunofluorescence remained in the peripheral region, and was significantly decreased (p<0.01) in the intracellular region. In the Stmn2-OE model, glucagon was localized mostly to the intracellular region along with Stmn2 **(Figure 2c, yellow arrowheads)**. Glucagon immunofluorescence intensity was significantly higher (p<0.0001) in the intracellular region and significantly decreased (p<0.0001) in the peripheral region upon Stmn2 overexpression (**Figure 2e**).

**Figure 2.**
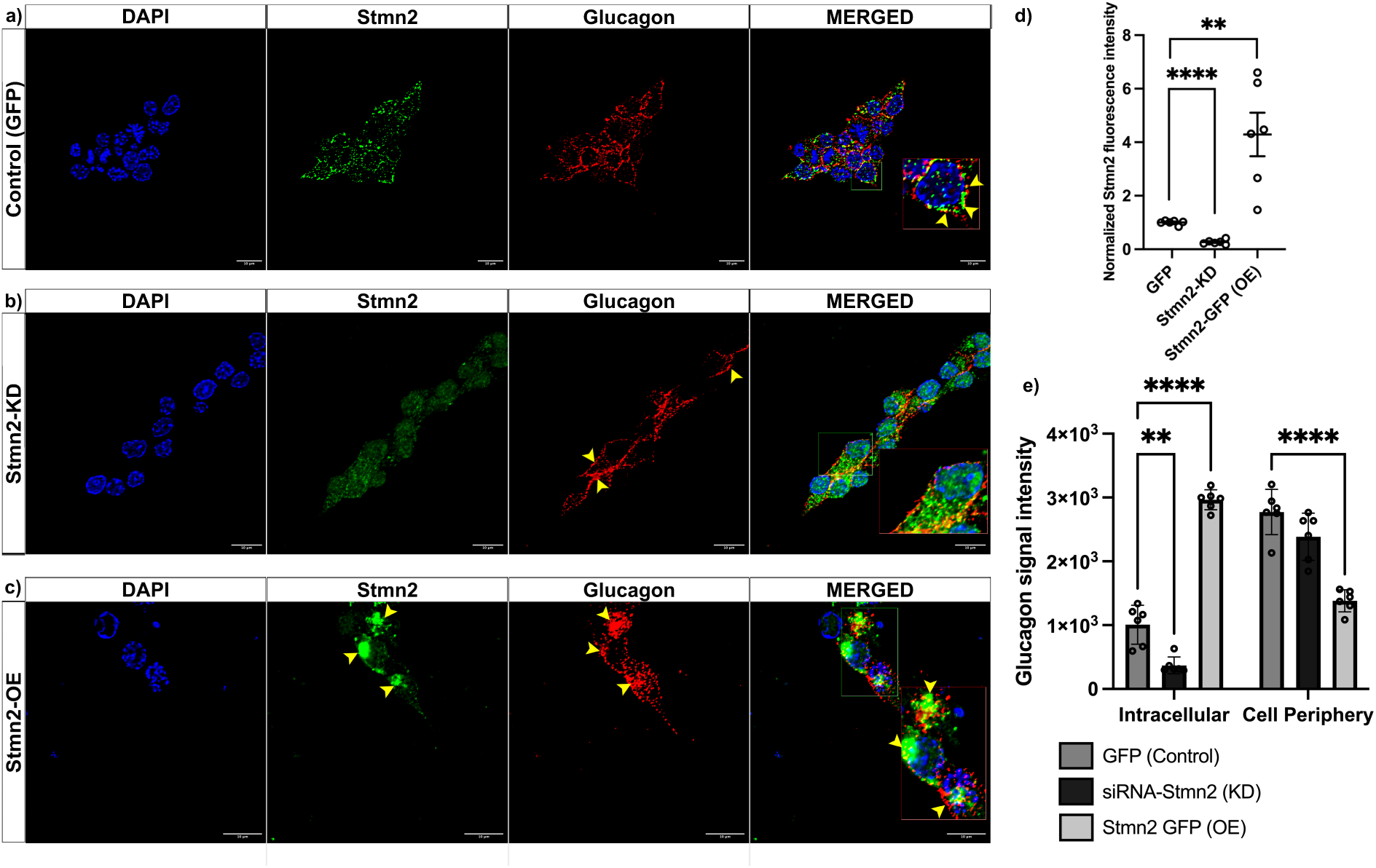
Glucagon distribution changes in response to Stmn2 silencing (KD) Stmn2-GFP (OE). Transfected cells were immunostained using primary antibodies against Stmn2 (green) and glucagon (red). **a)** Cells transfected with GFP alone show co-localized signals mainly at the cell periphery (yellow arrowheads). **b)** In Stmn2-KD cells, there is diminished Stmn2 immunofluorescence with glucagon localized largely to the periphery (yellow arrowheads). **c)** After Stmn2 OE, there is increased Stmn2 fluorescence intensity colocalizing strongly with glucagon in the intracellular region (yellow arrowheads). **d)** Changes in integrated Stmn2 fluorescence intensity in the Stmn2-KD and OE compared to the control (GFP-alone), normalized to endogenous Stmn2. Values were expressed as average integrated fluorescence intensity ±SEM (n=6) and compared among groups using a one-way ANOVA, followed by a post-hoc test. **p<0.01, ****p<0.0001. **e)** Glucagon distribution under control conditions (dark grey bars), and in response to Stmn2-KD (black bars) or OE (light grey bars) with values expressed as average glucagon fluorescence intensity ±SEM (n=6), compared among groups and cellular regions using a two-way ANOVA, followed by a post-hoc test. **p<0.01, ****p<0.0001.

To determine if Stmn2 was required for insulin-mediated glucagon trafficking, we treated αTC1-6 cells with 1 nM insulin after Stmn2 silencing or overexpression (**Figure 3**). In cells transfected with GFP alone and treated with insulin, glucagon distribution was mainly in the intracellular region (yellow arrowhead) (**Figure 3a**). In contrast, after Stmn2 KD, glucagon immunofluorescence remained largely in the peripheral region after treatment with insulin (**Figure 3b**, yellow arrowheads). In cells overexpressing Stmn2, glucagon immunofluorescence was again largely in the intracellular region after insulin treatment (**Figure 3c**, yellow arrowheads). Plot profile analysis **(Figure 3d)** showed that, in control cells treated with 1 nM insulin, glucagon was distributed largely to the intracellular region. After Stmn2 silencing, the effect of insulin was lost, as glucagon immunofluorescence in the intracellular region significantly decreased (p<0.0001) and significantly increased (p<0.0001) in the periphery. After Stmn2 OE, the distribution pattern of glucagon did not differ from that of insulin treatment in control cells (**Figure 3d**).

**Figure 3.**
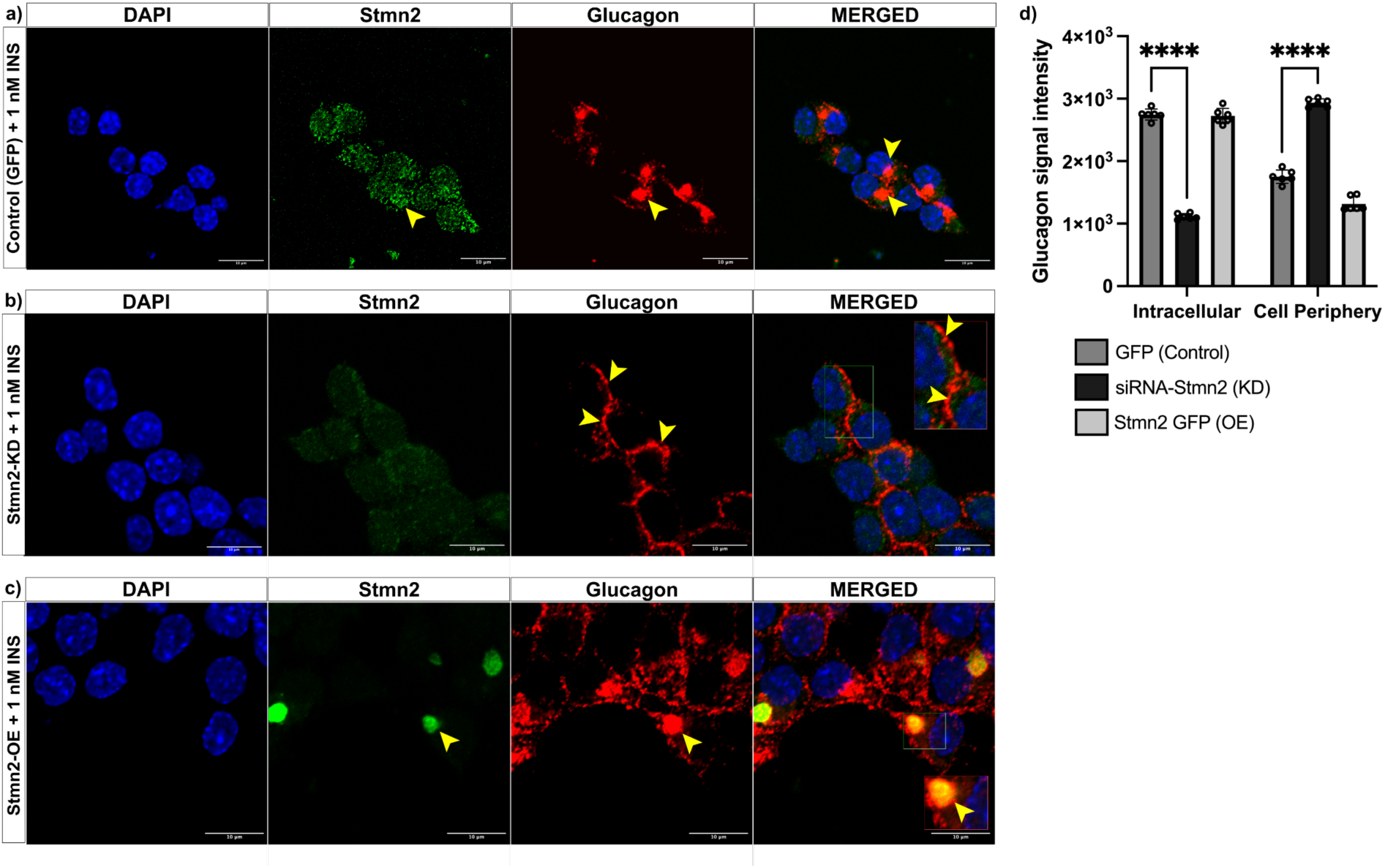
Glucagon distribution changes in the presence of 1 nM insulin in cells with Stmn2-GFP (OE) and Stmn2 siRNAs (KD). Transfected cells were treated with 1 nM insulin for 24 hours, then immunostained using primary antibodies against Stmn2 and glucagon. **a)** Cells transfected with GFP vector and treated with 1 nM insulin showed Stmn2 (green) and glucagon (red) redistributing intracellularly. **b)** The Stmn2-KD model diminished Stmn2 fluorescence and resulted in glucagon in the periphery, even after 1 nM insulin treatment. **c)** The Stmn2-OE model increased Stmn2 fluorescence intensity that colocalized strongly with glucagon in the intracellular region after 1 nM insulin treatment. **d)** Quantification of glucagon distribution in response to Stmn2-KD and OE with values expressed as average glucagon fluorescence intensity ±SEM (n=6), compared among groups and cellular regions using a two-way ANOVA, followed by a post-hoc test. ****p<0.0001.

To determine if Stmn2 was required for somatostatin-mediated glucagon trafficking, we treated aTC1-6 cells with 400 nM somatostatin after Stmn2 silencing or overexpression (**Figure 4**). In control cells transfected with GFP alone, glucagon distribution was mainly in the intracellular region (yellow arrowhead) (**Figure 4a**). In contrast, after Stmn KD, glucagon immunofluorescence appeared to be largely in the peripheral region after treatment with somatostatin (**Figure 4b**, yellow arrowheads). In cells overexpressing Stmn2, glucagon immunofluorescence was again largely in the intracellular region after somatostatin treatment (**Figure 4c**, yellow arrowheads). Plot profile analysis **(Figure 4d)** showed that, in control cells treated with 400 nM somatostatin, glucagon was distributed largely to the intracellular region. After Stmn2 silencing, the effect of somatostatin was lost, as glucagon immunofluorescence in the intracellular region significantly decreased (p<0.0001) and significantly increased (p<0.0001) in the periphery. After Stmn2 OE, the distribution of glucagon to the intracellular region did not differ from that of somatostatin treatment in control cells (Figure 4d). However, there was a significant decrease (p<0.0001) in glucagon in the peripheral region in somatostatin-treated cells compared with control cells (**Figure 4d**).

**Figure 4.**
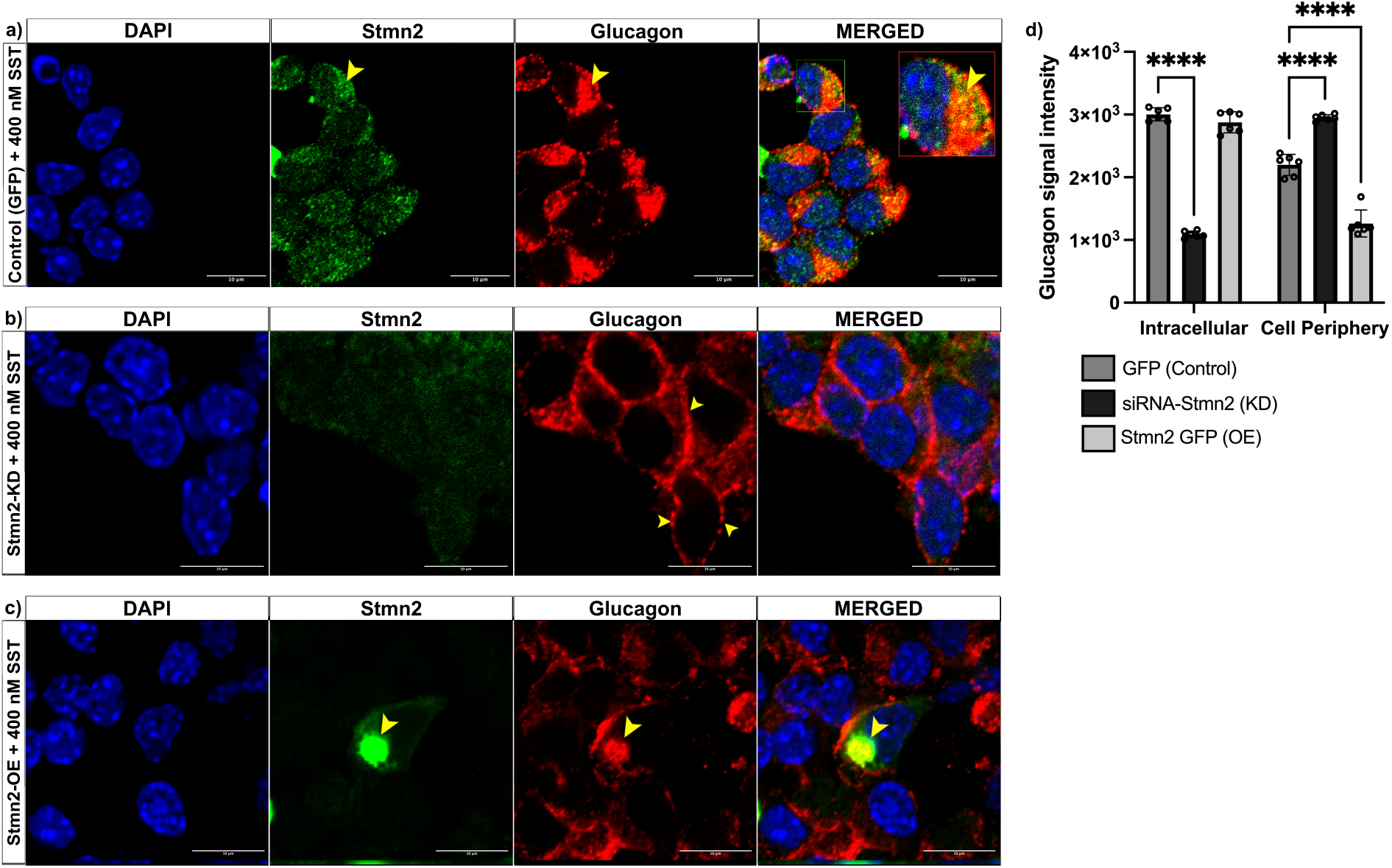
Fixed-cell immunostaining showing glucagon distribution changes in the presence of 400 nM somatostatin in cells with Stmn2-GFP (OE) and Stmn2 siRNAs (KD). Transfected cells were treated with 400 nM somatostatin for 24 hours. Cells were then immunostained using primary antibodies against Stmn2 (green) and glucagon (red). **a)** Cells transfected with GFP vector and treated with 400 nM somatostatin resulted in Stmn2 and glucagon distributing mostly in the intracellular region (yellow arrowheads). **b)** The Stmn2-KD model diminished Stmn2 fluorescence intensity, and glucagon was present in the periphery after somatostatin treatment. **c)** The Stmn2-OE model increased Stmn2 fluorescence intensity while colocalizing strongly with glucagon in the intracellular region after somatostatin treatment. **d)** Glucagon distribution in response to Stmn2-KD (red bars) and OE (green bars) with values expressed as average glucagon fluorescence intensity ±SEM (n=6) and compared among groups and cellular regions using a two-way ANOVA, followed by a post-hoc test. ****p<0.0001.

### Stmn2 mediates lysosomal dynamics in alpha cells

In order to visualize the spatial dynamics of lysosomes as regulated by Stmn2, we performed experiments utilizing cells transfected with the lysosomal fluorescent reporter LAMP1-RFP together with Stmn2-GFP (overexpression) or siRNA Stmn2 (silencing). In control cells expressing the GFP vector alone, LAMP1-RFP was distributed in the cell periphery and in the intracellular region (**Figure 5a, top row**). Stmn2-GFP overexpression caused almost complete colocalization with LAMP1-RFP in the intracellular region, with almost no fluorescence in the cell periphery **(Figure 5a, bottom row)**. Similar to the GFP vector alone, cells transfected with scrambled siRNAs displayed LAMP1-RFP signals in the cell periphery and some in the intracellular region **(Figure 5b, top row)**. In contrast, siRNA-mediated Stmn2 knockdown (KD) resulted in LAMP1-RFP localizing exclusively to the cell periphery **(Figure 5b, bottom row)**.

**Figure 5.**
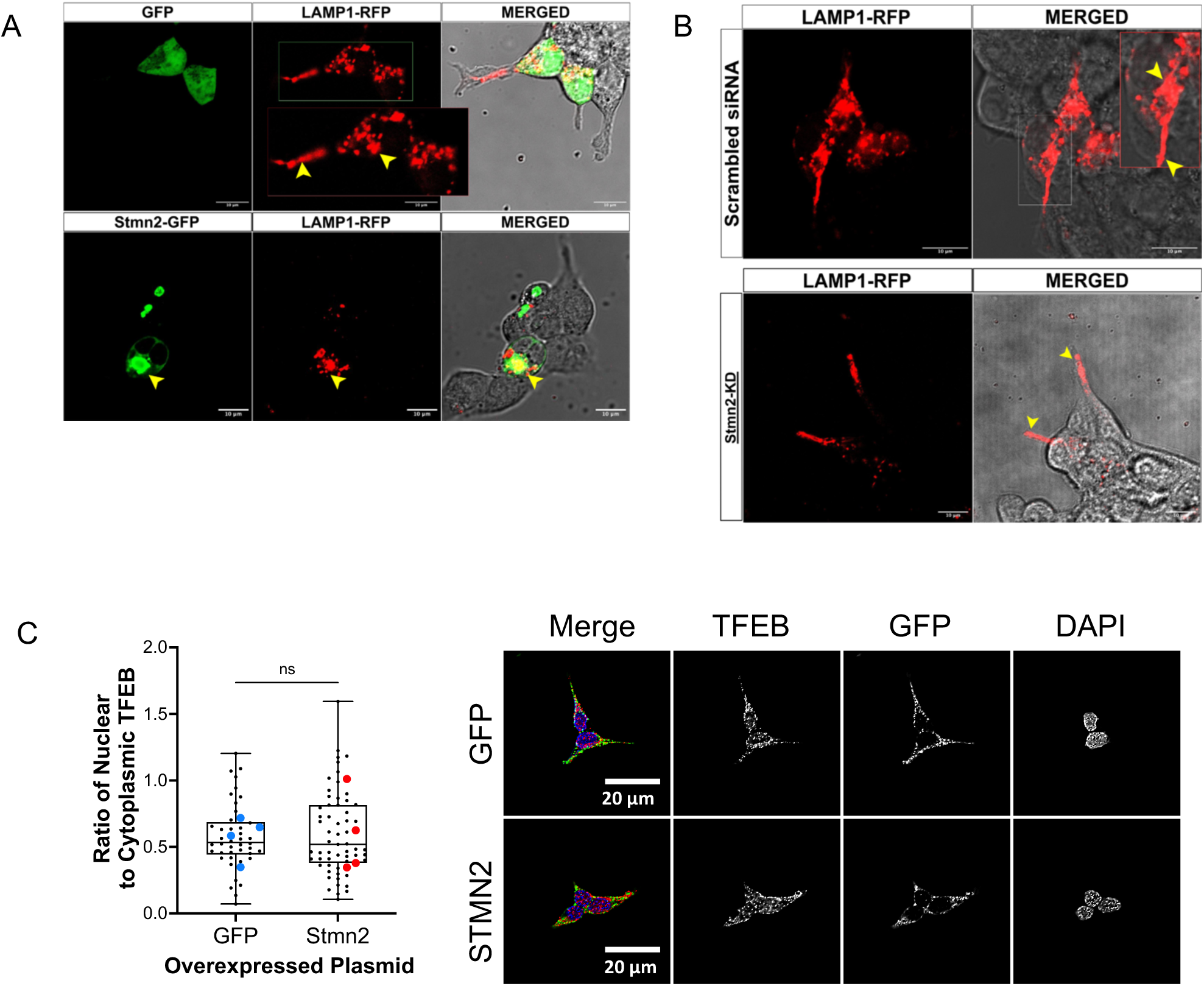
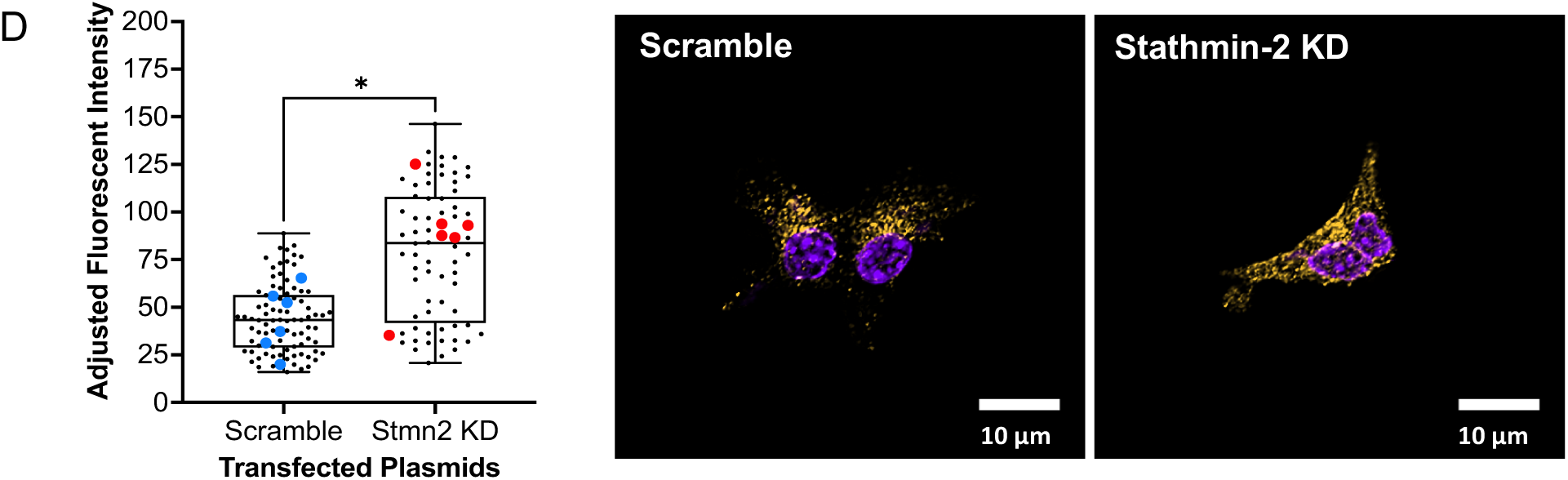
Stmn2 regulates lysosomal dynamics in αTC1-6 cells. Cells (n=3) were co-transfected with LAMP1-RFP and GFP vector or Stmn2-GFP, or with scrambled siRNA or siRNA against stathmin-2 (Stmn2-KD) and live images were captured 24 hours post-transfection. **a)** LAMP1-RFP is present in the cell periphery and the intracellular region (yellow arrowheads) after co-transfection with GFP alone as a negative control (upper panel). After co-transfection with Stmn2-GFP, (lower panel), LAMP1-RFP appears exclusively in the intracellular region (yellow arrowheads). **b)** LAMP1-RFP is present in the cell periphery and the intracellular region (yellow arrowheads) after co-transfection with scrambled siRNA sequences as a negative control for Stmn2-KD (upper panel). Transfection with Stmn2 siRNAs (KD) resulted in LAMP1-RFP exclusively in the cell periphery (yellow arrowheads) (lower panel). **c)** Transcription factor EB (TFEB) is not translocated to the nucleus upon overexpression of Stmn2-GFP. Immunofluorescence images of fixed cells show the subcellular distribution of TFEB, GFP alone (upper panel), Stmn2-GFP (lower panel) and cell nuclei (DAPI). Box plots show the ratio of nuclear to cytoplasmic TFEB immunofluorescence. Values were expressed as the average nuclear/cytoplasmic TFEB intensity ±SEM (n=4). Data points in colour represent the average ratio of 8-16 cells per coverslip; black points represent ratios from individual cells. **d)** Stmn2 regulates the lysosomal transport protein Arl8. Immunofluorescence images of fixed cells show the presence of Arl8 (orange) in cells transfected with scrambled siRNA or siRNA against Stmn2. Quantification of fluorescence (left) shows that knockdown of Stmn2 (Stmn2-KD) significantly increased the fluorescence intensity of Arl8. Values are means ±SEM (n=6). Data points in colour represent the average fluorescence intensities in 8-16 cells per coverslip; black points represent values from individual cells.

Since these results and our previous work ^20,21^ indicated that Stmn2 may affect lysosomal dynamics in αTC1-6 cells, we wished to determine if Stmn2 increased lysosomal biogenesis through nuclear translocation of transcription factor EB (TFEB), a master regulator of lysosomal gene transcription ^27^. Cells were transfected with GFP alone or Stmn2-GFP, and immunofluorescence distribution of TFEB was determined (**Figure 5c, yellow arrowheads**). Quantification of the nuclear:cytoplasmic ratio (**Figure 5c)** showed that overexpression of Stmn2 had no effect on the nuclear TFEB signal intensity.

We then investigated a possible role for Stmn2 in microtubule transport of lysosomes. Knockdown of Stmn2 in αTC1-6 cells increased immunofluorescence of Arl8, a small GTPase that functions in lysosomal transport along microtubules ^28^ (**Figure 5d**). Taken together, our results suggest that Stmn2 may mediate lysosomal dynamics in alpha cells through lysosomal transport and positioning, and not through transcriptional mechanisms.

### Insulin and somatostatin regulate lysosomal positioning in a Stmn2-dependent manner

Upon observing that the intracellular localization of glucagon+/LAMP1+ vesicles can be regulated by insulin or SST (**Figure 1**), and that Stmn2 may regulate lysosomal positioning (**Figure 5**), we sought to determine if insulin and somatostatin altered lysosomal trafficking via a Stmn2-mediated mechanism. Live cell imaging and plot profile analysis show that, in response to 1 nM insulin, LAMP1-RFP fluorescence in the cell periphery was decreased while the fluorescence in the intracellular region increased in αTC1-6 cells transfected with scramble siRNA (**Figure 6a**). This insulin-induced change in lysosomal redistribution was captured frame-by-frame over the span of 15 seconds post-treatment **(Figure 7)**. In contrast, there was no insulin-induced redistribution of LAMP1-RFP fluorescence after Stmn2 knockdown **(Figure 6b)**.

**Figure 6.**
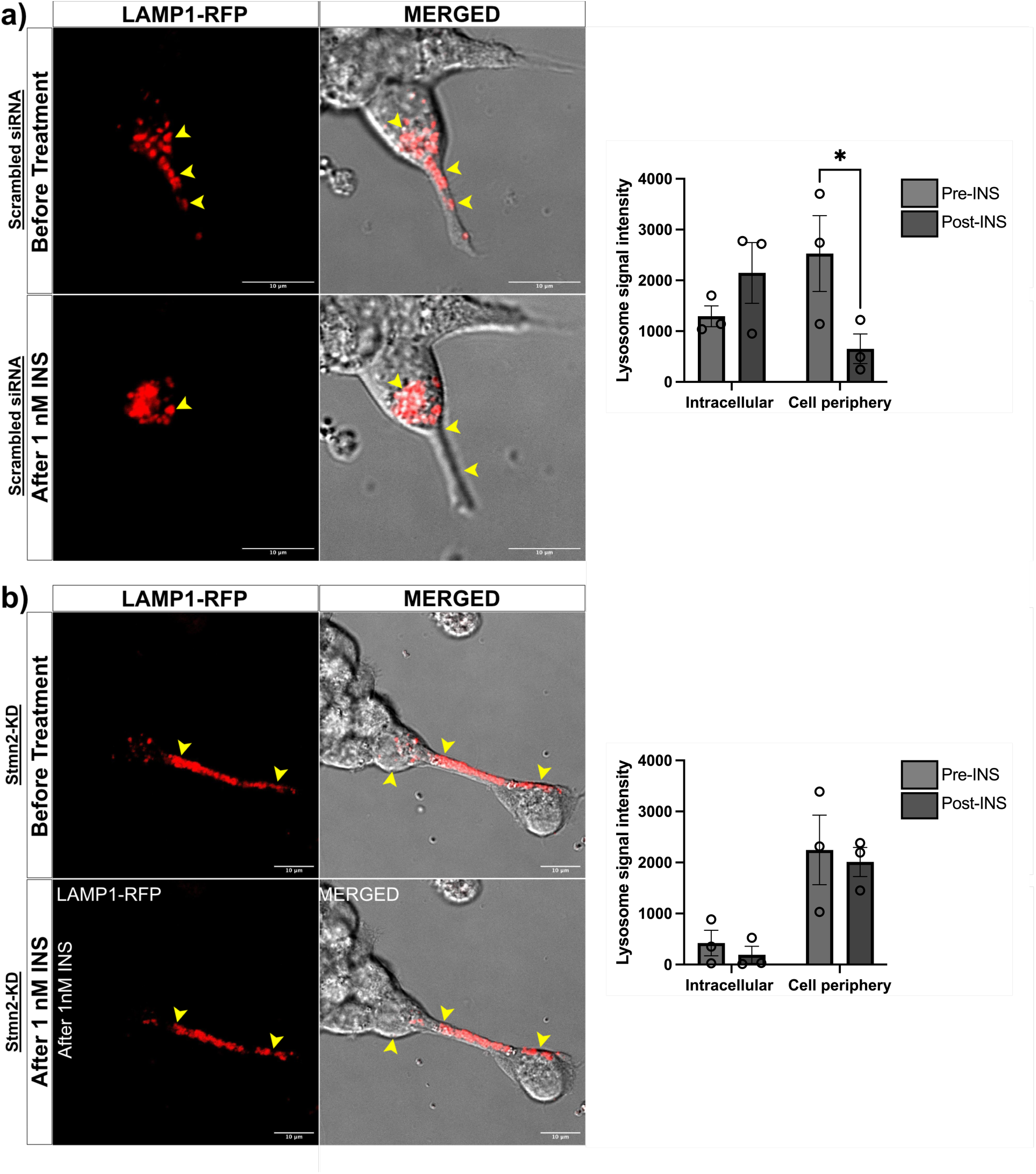
Stmn2 mediates the subcellular distribution of LAMP1-RFP in response to 1 nM insulin. Live images of αTC1-6 cells transfected with LAMP1-RFP (n=3) were captured in the same cell before, during, and five minutes after treatment with 1 nM insulin. **a)** LAMP1-RFP redistributed intracellularly (yellow arrowheads) in response to 1 nM insulin treatment in control cells transfected with scrambled siRNA. **b)** LAMP1-RFP distribution was not changed after 1 nM insulin treatment after siRNA-mediated knockdown of Stmn2 (Stmn2-KD) (yellow arrowheads). Graphs show the distribution of LAMP1-RFP fluorescence intensities in the intracellular and peripheral regions before (pre-INS) and after (post-INS) treatment with 1 nM insulin. Values are means ±SEM (n=3). * p<0.05

**Figure 7.**
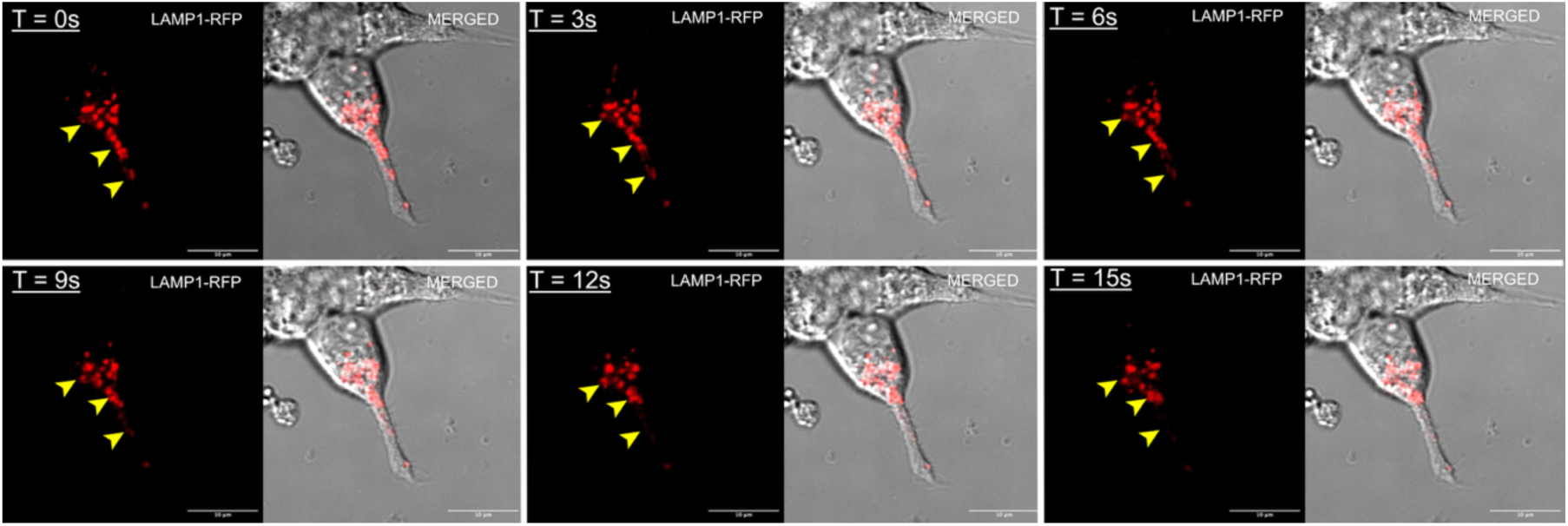
Live imaging of temporal changes in lysosomal dynamics in αTC1-6 cells upon 1 nM insulin treatment. Frame-by-frame capture (3s/ frame for 15s) of the image in Figure 3a) illustrates the immediate response of lysosomal redistribution in response to insulin (yellow arrowheads).

Treatment of control αTC1-6 cells with 400 nM somatostatin also elicited changes in lysosomal trafficking **(Figure 8a)** with a significant (p<0.01) redistribution to the intracellular region. Upon knockdown of Stmn2 **(Figure 8b)**, LAMP1-RFP remained in the cell periphery after treatment with somatostatin.

**Figure 8.**
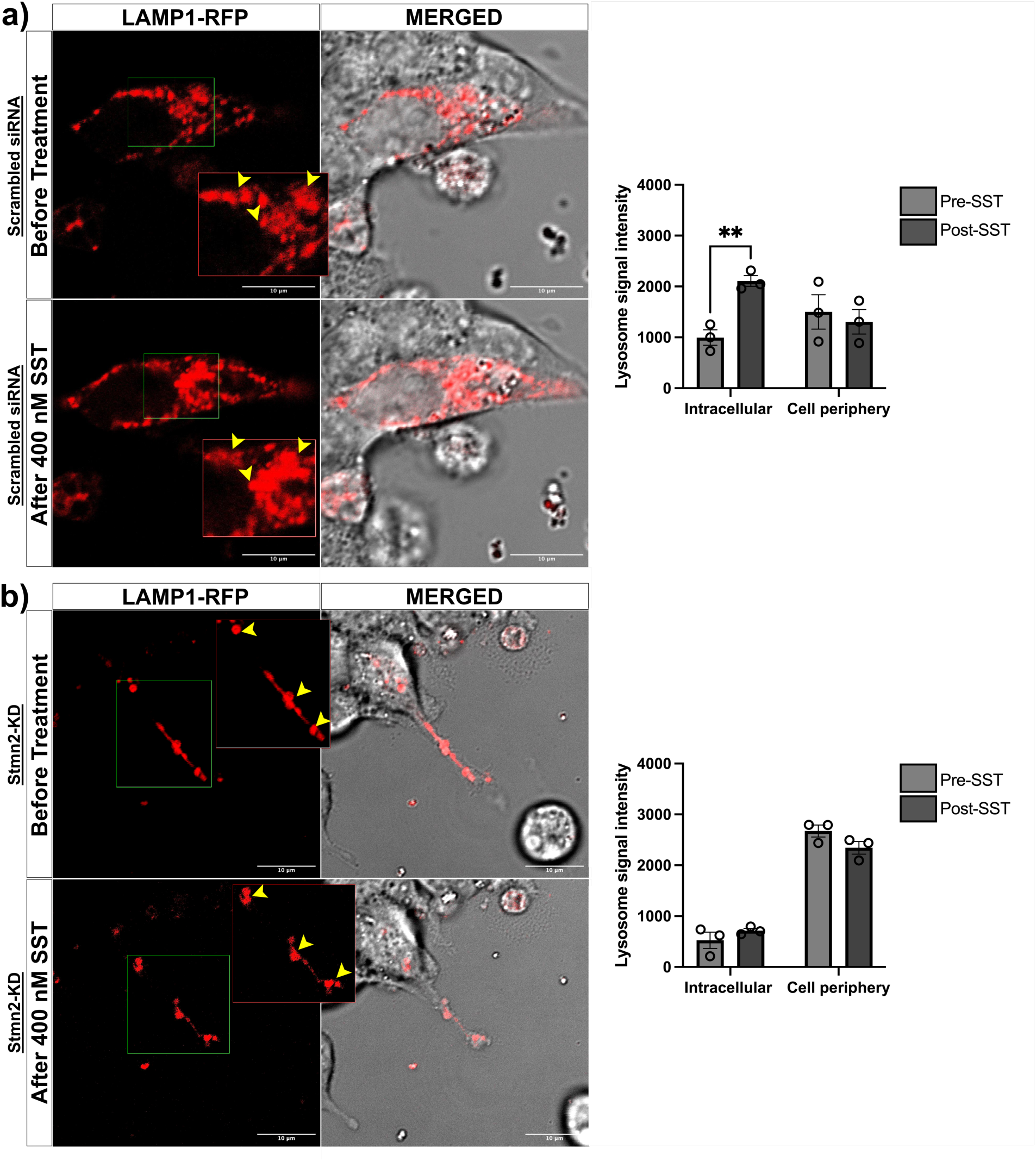
Stmn2 mediates the subcellular distribution of LAMP1-RFP in response to 400 nM somatostatin. αTC1-6 cells (n=3) were transfected overnight. Live images were captured in the same cell before, during, and five minutes after treatment with 400 nM somatostatin. **a)** LAMP1-RFP redistributed intracellularly (yellow arrowheads) in response to 400 nM somatostatin treatment in control cells transfected with scrambled siRNA. **b)** LAMP1-RFP distribution was not changed after 400 nM somatostatin treatment after siRNA-mediated knockdown of Stmn2 (Stmn2-KD) (yellow arrowheads). Graphs show means ±SEM (n=3). * p<0.05

## Discussion

Studies on the regulation of glucagon secretion have generally centered on secretory granule dynamics and exocytosis events at the plasma membrane. The role of intracellular transport mechanisms in the physiological regulation of glucagon in the pancreatic alpha cell is not well characterized. We have previously proposed that glucagon secretion may be regulated through lysosomal trafficking by the microtubule-binding protein, stathmin-2 (Stmn2) ^20^, and demonstrated that, in a mouse model of uncontrolled Type 1 diabetes, reduction in Stmn2 is associated with impaired excessive glucagon secretion and hyperglucagonemia ^21^. In this study, we investigated a role for Stmn2 in the physiological regulation of glucagon transport by the paracrine factors insulin and somatostatin.

The decrease of glucagon+/Stmn2+ vesicles present at the cell periphery upon treatment with either insulin+GABA or somatostatin suggests that these paracrine effectors may exert their inhibitory effects by relocating secretory vesicles away from sites of exocytosis. However, in dispersed human and mouse alpha cells, the number of secretory granules at sites of exocytosis was not affected by either insulin or somatostatin; instead, they acted by granule de-priming ^14,29^. As Lamp1+ lysosomes containing glucagon were significantly increased in the intracellular region by treatment with insulin + GABA or somatostatin, it is possible that the glucagon+/Stmn2+ vesicles were lysosomes. We have previously shown localization of Stmn2 in both secretory granules and lysosomes in αTC1-6 cells and in alpha cells in mouse islets ^20,21^, and BioID proximity proteomics has recently shown that Stmn2 may be present in the luminal fraction of secretory granules and lysosomes in neurons ^30^. Therefore, these results suggest that insulin and somatostatin facilitate the retrograde movement of glucagon contained within the lysosomal compartment, and not in secretory granules, away from the plasma membrane, and that there may be a role for Stmn2 in this transport mechanism. It has been suggested that Stmn2 plays a role in the targeting of some molecular cargo to the secretory granule ^31^, indicating that Stmn2 may function in the trafficking of glucagon to both secretory granules and lysosomes.

To examine the effects of insulin and somatostatin on glucagon at sites of exocytosis, we examined the localization of glucagon+/syntaxin1+ vesicles. Syntaxin 1A and SNAP-25 are t-SNARES that mediate exocytosis in alpha cells ^32,33^, and specifically, syntaxin 1A is transmembrane protein that associates with ion channels in primary mouse alpha cells and αTC1-6 cells at sites of glucagon granule exocytosis ^33^. The reduction by somatostatin of glucagon+/syntaxin1A+ vesicles suggests a reduction in the number of secretory granules at sites of exocytosis. However, there was also an increase in the intracellular region. As syntaxin 1A has not been localized to secretory granules, these results indicate that syntaxin1A associates with glucagon in another compartment. Recently, it has been shown that syntaxin 1A marks secretory lysosomes in non-neuronal cells ^34^. Taken together with the glucagon+/LAMP1 results, insulin and somatostatin may act on alpha cells to enhance the lysosomal trafficking of glucagon towards the intracellular region.

We next examined if insulin and somatostatin required Stmn2 for the intracellular routing of glucagon. We built upon our previous results ^20^ by showing that Stmn2 KD reduced intracellular glucagon, and overexpression of Stmn2 (OE) dramatically increased glucagon immunofluorescence in the intracellular region while decreasing peripherally-localized glucagon. Both insulin and somatostatin enhanced the intracellular localization of glucagon, and this effect was lost in the absence of Stmn2, with glucagon located largely at the cell periphery. These results indicate that insulin and somatostatin require the presence of Stmn2 for directing glucagon trafficking towards the intracellular region of αTC1-6 cells. While several studies have described how insulin receptor- and SSTR2-mediated signalling disrupt alpha cell secretory granule dynamics and membrane electrical activity ^14,29,35,36^, here we show that insulin and somatostatin may also redirect the intracellular trafficking of glucagon from the cell periphery to the intracellular regions in a Stmn2-dependent manner. Together with the results showing that insulin and somatostatin increased the distribution of glucagon+/LAMP1+ vesicles in the intracellular region, we reason that these paracrine factors regulate glucagon secretion by increasing the transport of glucagon+/Lamp1+ lysosomes to the perinuclear region in addition to inhibiting glucagon granule priming and exocytosis.

In order to more closely examine the mechanism underlying Stmn2-mediated intracellular transport in α cells, we first investigated the direct effects of Stmn2 on lysosomal transport in αTC1-6 cells. Overexpression of Stmn2 directed lysosomes, marked by LAMP1-RFP, to the intracellular region, while knockdown of Stmn2 resulted in dispersion of lysosomes to the cell periphery. As our previous results showed that overexpression of Stmn2 increased LAMP2 fluorescence intensity ^20^, we postulated that Stmn2 increased lysosomal biogenesis and function. The present study shows that Stmn2 does not appear to increase lysosomal biogenesis, as overexpression of Stmn2 did not result in nuclear translocation of TFEB, a master transcriptional regulator of lysosomal biogenesis and function ^27,37,38^. Instead, Stmn2 may regulate lysosomal transport in α cells in a mechanism involving the Arf-like small GTPase, Arl8 ^39^. The increase in Arl8 immunofluorescence in the absence of Stmn2 in αTC1-6 cells suggests a mechanism whereby Stmn2 directs the transport of lysosomes to the intracellular region by suppressing Arl8, and when Stmn2 is decreased, this inhibition is relieved, allowing Arl8 to function in the trafficking of lysosomes towards the cell periphery via kinesin-mediated anterograde transport along microtubules ^26,28^. It is well established that Stmn2 interacts with microtubules and binds to tubulin via tubulin-binding domains ^19,30^. Although the function of tubulin-bound Stmn2 is not known, we speculate that it may displace or interfere with the Arl8-HOPS-kinesin complex ^26,40,41^ to cause retrograde trafficking to the cell interior in alpha cells.

In contrast to its negative regulatory function in alpha cells, Stmn2 has been identified as essential for axonal regeneration and neural growth cone stabilization, and promotes lysosomal transport along the axon, possibly via kinesin-mediated anterograde transport ^42–44^. The absence of functional Stmn2 is associated with neurodegenerative diseases, particularly ALS and fronto-temporal dementia, in which axonal regeneration is inhibited, leading to neuronal damage that can be reversed by restoration of Stmn2 expression ^43,45,46^. Why Stmn2 would enhance anterograde transport of lysosomes in neurons and retrograde transport in alpha cells is not known, but we reason that these seemingly disparate functions maintain cell-specific homeostasis. Stmn2-mediated negative regulation of glucagon trafficking and secretion may be part of the normal physiology of the alpha cell, and enhanced basal secretion through lysosomes when Stmn2 is down-regulated may contribute to the pathophysiology of the hyperglucagonemia of diabetes, as we have previously suggested ^20–22,47^.

To directly determine if the effects of insulin and somatostatin on glucagon trafficking were through Stmn2-mediated retrograde lysosomal transport, αTC1-6 cells harbouring LAMP1-RFP were treated with insulin or somatostatin after siRNA-mediated knockdown of Stmn2. Insulin caused the rapid retrograde transport of lysosomes to the intracellular region in an Stmn2-dependent manner; to our knowledge, this action of insulin on alpha cells has not been previously documented. This interpretation is somewhat limited since we did not use an insulin receptor antagonist, although the insulin receptor is expressed on αTC1-6 cells ^23,48^. We speculate that insulin receptor signaling may alter the phosphorylation status of Stmn2 on lysosomes to then displace or override Arl8 and activate dynein-mediated retrograde transport ^27^. Such a mechanism requires further experimentation in primary alpha cells or isolated islets. Although we did not conduct a precise calculation of the velocity of this retrograde lysosomal movement, it appears that the LAMP1-RFP vesicles were moving within the velocity ranges of 1.38-1.74 μm/sec reported for the retrograde movement of fluorescently-tagged LAMP1 in hiPSC-derived cortical neurons ^49^.

Our data regarding the role of somatostatin in the Stmn2-mediated lysosomal trafficking of glucagon is less clear. That intracellular retention of glucagon, glucagon+/LAMP1+ vesicles and LAMP1-RFP occurs in response to somatostatin treatment aligns with the hypothesis that the alpha cell is under almost constant tonic inhibition by somatostatin; the alpha cell counter-regulatory response occurs at very low (0-1 mM) glucose, above which somatostatin secretion is stimulated to suppress glucagon secretion ^1,50^. However, unlike the effects of insulin, Stmn2 overexpression decreased glucagon at the cell periphery in the presence of somatostatin, and somatostatin treatment did not result in measurable retrograde movement of lysosomes from the periphery to the intracellular region of αTC1-6 cells. Therefore, somatostatin receptor signalling through SSTR2 may not play a significant role in Stmn2-mediated retrograde transport of lysosomes containing glucagon, but may figure in intracellular retention of lysosomes. There is evidence that G protein-coupled receptors can regulate cargo transport through interaction with microtubule machinery ^51^, and precisely how SSTR2-G_αi_-cAMP signalling in alpha cells regulate lysosomal dynamics remains to be determined.

Although our results may indicate a new pathway through which insulin and somatostatin negatively regulate glucagon secretion, the treatment of an alpha cell line with these hormones does not truly capture the paracrine mechanisms that regulate glucagon secretion within intact islets. Alpha, beta and delta cells operate in finely-tuned and multi-faceted feedback loops that involve intrinsic nutrient sensing and responses to paracrine inputs to regulate glucagon secretion ^1,22,52–54^. For this reason, further investigations of Stmn2-mediated lysosomal regulation of glucagon trafficking by paracrine factors need to be conducted *in vivo*, in pancreatic slices, or in isolated islets, with functional assays of glucagon secretion. Nonetheless, we and others have been able to recapitulate findings in αTC1-6 cells in mouse islets and rat models ^21,55,56^, and there is a report of altered lysosomal trafficking of glucagon in human alpha cells from T1D donors ^57^, providing evidence that αTC1-6 cells are a useful tool for initial investigations that lead to novel insights in *in vivo* animal models and human-derived cells and tissues.

In conclusion, our data suggest that insulin and somatostatin may inhibit glucagon secretion through regulating lysosomal trafficking of glucagon in addition to regulating secretory granule dynamics. We propose that insulin-mediated inhibition of glucagon may occur through Stmn2-mediated retrograde trafficking of glucagon-containing lysosomes in a mechanism involving modulation of the small GTPase Arl8 that regulates lysosomal positioning.

## Acknowledgements

This work was supported by a Natural Sciences and Engineering Research Council of Canada (NSERC) Discovery Grant to SD. NC was supported by a scholarship from the Canadian Islet Research and Training Network (CIRTN-R2FIC) funded by NSERC-CREATE.

## References

1. Noguchi, G. M. & Huising, M. O. Integrating the inputs that shape pancreatic islet hormone release. Nat. Metab. 1, 1189–1201 (2019).

2. Brereton, M. F., Vergari, E., Zhang, Q. & Clark, A. Alpha-, Delta- and PP-cells. Journal of Histochemistry & Cytochemistry 63, 575–591 (2015).

3. Hædersdal, S., Andersen, A., Knop, F. K. & Vilsbøll, T. Revisiting the role of glucagon in health, diabetes mellitus and other metabolic diseases. Nat. Rev. Endocrinol. 19, 321–335 (2023).

4. Ma, X. et al. Glucagon Stimulates Exocytosis in Mouse and Rat Pancreatic α-Cells by Binding to Glucagon Receptors. Molecular Endocrinology 19, 198–212 (2005).

5. Leibiger, B. et al. Glucagon regulates its own synthesis by autocrine signaling. Proc. Natl. Acad. Sci. U. S. A. 109, 20925–30 (2012).

6. Gromada, J., Franklin, I. & Wollheim, C. B. Alpha-cells of the endocrine pancreas: 35 years of research but the enigma remains. Endocr. Rev. 28, 84–116 (2007).

7. Wendt, A. et al. Glucose Inhibition of Glucagon Secretion From Rat {a}-Cells Is Mediated by GABA Released From Neighboring {b} cells. Diabetes 53, 1038–1045 (2004).

8. Li, C. et al. Regulation of glucagon secretion in normal and diabetic human islets by ??-hydroxybutyrate and glycine. Journal of Biological Chemistry 288, 3938–3951 (2013).

9. Rorsman P, Berggren PO, Bokvist K, Ericson H, Möhler H, Ostenson CG, S. P. Glucose-inhibition of glucagon secretion involves activation of GABAA-receptor chloride channels. Nature 341, 233–236 (1989).

10. Rutter, G. A. Regulating Glucagon Secretion: Somatostatin in the Spotlight. Diabetes 58, 299–301 (2009).

11. Xu, S. F. S., Andersen, D. B., Izarzugaza, J. M. G., Kuhre, R. E. & Holst, J. J. In the rat pancreas, somatostatin tonically inhibits glucagon secretion and is required for glucose-induced inhibition of glucagon secretion. Acta Physiologica 229, 1–15 (2020).

12. Briant, L. J. B. et al. δ-cells and β-cells are electrically coupled and regulate α-cell activity via somatostatin. J. Physiol. 596, 197–215 (2018).

13. Elliott, A. D., Ustione, A. & Piston, D. W. Somatostatin and insulin mediate glucose-inhibited glucagon secretion in the pancreatic α-cell by lowering cAMP. Am. J. Physiol. Endocrinol. Metab. 308, E130–E143 (2015).

14. Gromada, J., Hoy, M., Bushcard, K., Salehi, A. & Rorsman, P. Somatostatin inhibits exocytosis in rat pancreatic a-cells by Gi2-dependent activation of calcineurin and depriming of secretory granules. Journal of Physiology 535, 519–532 (2001).

15. Vergari, E. et al. Insulin inhibits glucagon release by SGLT2-induced stimulation of somatostatin secretion. Nat. Commun. 10, 139 (2019).

16. Goginashvili, A. et al. Insulin secretory granules control autophagy in pancreatic β cells. Science (1979). 347, 878–882 (2015).

17. Riahi, Y. et al. Autophagy is a major regulator of beta cell insulin homeostasis. Diabetologia 59, 1480–1491 (2016).

18. Rajak, S. et al. MTORC1 inhibition drives crinophagic degradation of glucagon. Mol. Metab. 53, (2021).

19. Chauvin, S. & Sobel, A. Neuronal stathmins: A family of phosphoproteins cooperating for neuronal development, plasticity and regeneration. Progress in Neurobiology vol. 126 1–18 Preprint at 10.1016/j.pneurobio.2014.09.002 (2015).

20. Asadi, F. & Dhanvantari, S. Stathmin-2 Mediates Glucagon Secretion From Pancreatic α-Cells. Front. Endocrinol. (Lausanne). 11, 1–13 (2020).

21. Asadi, F. & Dhanvantari, S. Misrouting of glucagon and stathmin-2 towards lysosomal system of α-cells in glucagon hypersecretion of diabetes. Islets 14, 40–57 (2022).

22. Asadi, F. & Dhanvantari, S. Pathways of Glucagon Secretion and Trafficking in the Pancreatic Alpha Cell: Novel Pathways, Proteins, and Targets for Hyperglucagonemia. Front. Endocrinol. (Lausanne). 12, (2021).

23. Asadi, F. & Dhanvantari, S. Plasticity in the glucagon interactome reveals novel proteins that regulate glucagon secretion in α-TC1-6 cells. Front. Endocrinol. (Lausanne). 10, 792 (2019).

24. Kumar, G. et al. RUFY3 links Arl8b and JIP4-Dynein complex to regulate lysosome size and positioning. Nat. Commun. 13, 1–21 (2022).

25. Jongsma, M. L. L. M. et al. An ER-Associated Pathway Defines Endosomal Architecture for Controlled Cargo Transport. Cell 166, 152–166 (2016).

26. Guardia, C. M., Farías, G. G., Jia, R., Pu, J. & Bonifacino, J. S. BORC Functions Upstream of Kinesins 1 and 3 to Coordinate Regional Movement of Lysosomes along Different Microtubule Tracks. Cell Rep. 17, 1950–1961 (2016).

27. Settembre, C. & Perera, R. M. Lysosomes as coordinators of cellular catabolism, metabolic signalling and organ physiology. Nat. Rev. Mol. Cell Biol. 25, 223–245 (2024).

28. Rosa-Ferreira, C. & Munro, S. Arl8 and SKIP Act Together to Link Lysosomes to Kinesin-1. Dev. Cell 21, 1171–1178 (2011).

29. Omar-Hmeadi, M., Lund, P. E., Gandasi, N. R., Tengholm, A. & Barg, S. Paracrine control of α-cell glucagon exocytosis is compromised in human type-2 diabetes. Nat. Commun. 11, 1–11 (2020).

30. Deng, X., Bradshaw, G. A., Kalocsay, M. & Mitchison, T. Tubulin regulates stability and localization of STMN2 by binding preferentially to its soluble form. J. Cell Biol. 224, e202502192 (2025).

31. Mahapatra, N. R., Taupenot, L., Courel, M., Mahata, S. K. & O’Connor, D. T. The trans-golgi proteins SCLIP and SCG10 interact with chromogranin A to regulate neuroendocrine secretion. Biochemistry 47, 7167–7178 (2008).

32. Andersson, S. A., Pedersen, M. G., Vikman, J. & Eliasson, L. Glucose-dependent docking and SNARE protein-mediated exocytosis in mouse pancreatic alpha-cell. Pflugers Arch. 462, 443–454 (2011).

33. Xia, F. et al. Targeting of voltage-gated K+ and Ca2+ channels and soluble N-ethylmaleimide-sensitive factor attachment protein receptor proteins to cholesterol-rich lipid rafts in pancreatic α-cells: Effects on glucagon stimulus-secretion coupling. Endocrinology 148, 2157–2167 (2007).

34. Bhatt, A. M., Singh, B., Singh, P. & Setty, S. R. G. STX1A localizes to the lysosome and controls its exocytosis. Mol. Biol. Cell 36, ar153 (2025).

35. Leung, Y. M. et al. Insulin regulates islet alpha-cell function by reducing KATP channel sensitivity to adenosine 5’-triphosphate inhibition. Endocrinology 147, 2155–2162 (2006).

36. Kawamori, D. & Kulkarni, R. N. Insulin modulation of glucagon secretion: the role of insulin and other factors in the regulation of glucagon secretion. Islets 1, 276–279 (2009).

37. Sardiello, M. et al. A gene network regulating lysosomal biogenesis and function. Science (1979). 325, 473–477 (2009).

38. Palmieri, M. et al. Characterization of the CLEAR network reveals an integrated control of cellular clearance pathways. Hum. Mol. Genet. 20, 3852–3866 (2011).

39. Khatter, D., Sindhwani, A. & Sharma, M. Arf-like GTPase Arl8: Moving from the periphery to the center of lysosomal biology. Cell. Logist. 5, e1086501 (2015).

40. Hummel, J. J. A. & Hoogenraad, C. C. Specific kif1a–adaptor interactions control selective cargo recognition. Journal of Cell Biology 220, (2021).

41. Shelke, G. V., Williamson, C. D., Jarnik, M. & Bonifacino, J. S. Inhibition of endolysosome fusion increases exosome secretion. Journal of Cell Biology 222, (2023).

42. Lutjens, R. et al. Localization and targeting of SCG10 to the trans-Golgi apparatus and growth cone vesicles. European Journal of Neuroscience 12, 2224–2234 (2000).

43. Baughn, M. W. et al. Mechanism of STMN2 cryptic splice-polyadenylation and its correction for TDP-43 proteinopathies. Science (1979). 379, 1140–1149 (2023).

44. Tymanskyj, S. R., Curran, B. M. & Ma, L. Selective axonal transport through branch junctions is directed by growth cone signaling and mediated by KIF1/kinesin-3 motors. Cell Rep. 39, 110748 (2022).

45. Prudencio, M. et al. Truncated stathmin-2 is a marker of TDP-43 pathology in frontotemporal dementia. Journal of Clinical Investigation 130, 6080–6092 (2020).

46. Melamed, Z. et al. Premature polyadenylation-mediated loss of stathmin-2 is a hallmark of TDP-43-dependent neurodegeneration. Nat. Neurosci. 22, 180–190 (2019).

47. Dhanvantari, S. & Dean, E. D. Glucagon, the Alpha Cell, and Potential Targets for Diabetes Treatment. Endocrinology (United States) 166, bqaf162 (2025).

48. McGirr, R. et al. Glucose Dependence of the Regulated Secretory Pathway in {a}TC1-6 Cells. Endocrinology 146, 4514–4523 (2005).

49. Abouward, R. et al. LAMP1 and LAMP2A localise to axonal organelles with distinct motility dynamics and partially overlapping molecular signatures in human neurons. J Cell Science jcs264466, (2026).

50. Gao, R. et al. Antecedent hypoglycaemia impairs glucagon secretion by enhancing somatostatin-mediated negative feedback control. Nat. Metab. https://doi.org/10.1038/s42255-025-01422-7 (2026) doi:10.1038/s42255-025-01422-7.

51. Schappi, J. M., Krbanjevic, A. & Rasenick, M. M. Tubulin, actin and heterotrimeric G proteins: Coordination of signaling and structure. Biochim. Biophys. Acta Biomembr. 1838, 674–681 (2014).

52. Watts, M., Ha, J., Kimchi, O. & Sherman, A. Paracrine regulation of glucagon secretion: the β/α/δ model. American Journal of Physiology-Endocrinology and Metabolism 310, E597–E611 (2016).

53. Gao, R. et al. α-cell electrophysiology and the regulation of glucagon secretion. J. Endocrinol. 258, e220295 (2023).

54. Ng, X. W., Kong, C., DiGruccio, M. R., Lee, J. & Piston, D. W. Role of complexin 2 in the regulation of hormone secretion from the islet of Langerhans. Am. J. Physiol. Endocrinol. Metab. 329, E861–E873 (2025).

55. Diao, J., Asghar, Z., Chan, C. B. & Wheeler, M. B. Glucose-regulated glucagon secretion requires insulin receptor expression in pancreatic alpha-cells. J. Biol. Chem. 280, 33487–96 (2005).

56. Asadi, F. et al. Sex-dependent Effect of In-utero Exposure to Δ9-Tetrahydrocannabinol on Glucagon and Stathmin-2 in Adult Rat Offspring. Can. J. Diabetes 46, 851–862 (2022).

57. dos Santos, T., et al. Altered immune and metabolic molecular pathways drive islet cell dysfunction in human type 1 diabetes. Journal of Clinical Investigation 135, 1–17 (2025).

